# Improving the efficacy and accessibility of intracranial viral vector delivery in non-human primates

**DOI:** 10.1101/2022.06.06.494543

**Authors:** Devon J Griggs, Aaron D. Garcia, Wing Yun Au, William K. S. Ojemann, Andrew Graham Johnson, Jonathan T. Ting, Elizabeth A. Buffalo, Azadeh Yazdan-Shahmorad

## Abstract

Non-human primates (NHPs) are precious resources for cutting edge neuroscientific research, including large-scale viral vector-based experimentation such as optogenetics. We propose to improve surgical outcomes by enhancing surgical preparation practices of convection-enhanced delivery (CED), which is an efficient viral vector infusion technique for large brains such as NHPs’. Here we present both real-time and next-day MRI data of CED in the brains of ten NHPs, and we present a quantitative, inexpensive, and practical bench-side model of the *in vivo* CED data. Our bench-side model is composed of food coloring infused into a transparent agar phantom, and the spread of infusion is optically monitored over time. Our proposed method approximates CED infusions into the cortex, thalamus, medial temporal lobe and caudate nucleus of NHPs, confirmed by MRI data acquired with either gadolinium-based or manganese-based contrast agents co-infused with optogenetic viral vectors. These methods and data serve to guide researchers and surgical team members in key surgical preparations for intracranial viral delivery using CED in NHPs, and thus improve expression targeting and efficacy and, as a result, reduce surgical risks.

## 1. Introduction

The development of novel tools to genetically alter the properties of neurons has been instrumental in expanding the scope of neuroscientific questions. Perhaps one of the most popular of these genetic modification methods is optogenetics, through which cells can be made susceptible to rapid, reversible manipulations via light stimulation. This technique was first demonstrated in 2005 [1] and since then has been robustly developed and widely adopted. In particular, optogenetics has become a powerful tool for rodent neuroscience [2], [3]. The high-throughput nature of experiments with these models means that researchers can pilot, experiment, and make corrections with new subjects rapidly and with minimal resources. Additionally, the rapid gestational time (∼1 month for most research species) and large litter size (4-12 pups/litter) of these models extends the genetic modification toolkit of rodent researchers to allow for breeding genetically altered lines and testing transgenic subjects with relative ease. However, transgenic lines are largely unavailable for highly translational non-human primate (NHPs) models. In contrast to rodents, the macaque has a gestational period of ∼6 months and gives birth to singular offspring, which eliminates the practical viability of transgenic approaches. This leaves viral vector transfection as the primary method of preparing NHPs for experiments requiring genetic modification. However, this experimental approach still proves to be a challenging, and potentially costly, endeavor. Here, we present novel and updated methods that eliminate several of the main dissuading factors of transduction studies in NHPs.

Convection enhanced delivery (CED) is an infusion technique which has been developed over the past few decades to deliver medicinal agents to the brain [4], [5]. Classic neural infusion techniques rely on diffusion which is based on a concentration gradient and therefore is heavily influenced by molecular weight [6]. Large molecules like viral vectors are inefficiently spread by diffusion. By contrast, CED capitalizes on a pressure gradient for delivery which is much less influenced by molecular weight and comes with a number of benefits over diffusion: 1-CED can be performed at higher delivery rates (on the order of 1 µL/min or higher) which speeds up delivery [7]–[9], 2-CED can distribute agent over greater volumes (hundreds of mm^3^) [8], [9], 3-CED produces a roughly uniform concentration of agent throughout the distribution volume [6], [7], [10]–[14], and 4-as previously alluded, CED is effective at transporting large molecules, such as viral vectors, throughout regions of the brain [7], [11]–[15].

Incentivized by these desirable properties, we have used CED in recent years to deliver optogenetic viral vectors into the brains of non-human primates for neuroscientific experiments [8], [9], [16], [17]. To compliment the technique, we have utilized a method of real-time infusion validation with magnetic resonance imaging (MRI) technology using a gadolinium-based contrast agent co-infused with the optogenetic viral vector [18]. We previously validated the resulting optogenetic expression with epifluorescence imaging, electrophysiology, and histology [8], [9]. However, we recognize that not all institutions have access to an MRI scanner in which live validation of an injection can be visualized. Thus, a separate set of CED experiments without live MRI guidance was performed using novel optogenetic viral vectors co-infused with a manganese-based contrast agent. This allowed us to confirm infusion success the following day with MRI. With these live and next-day *in vivo* MRI data collected across ten animals and three different brain regions, we were uniquely positioned to develop a model to assist in planning CED procedures in the brains of NHPs. This work is important to the field because not all researchers have access to MRI scanners for next-day imaging, and fewer still are equipped to perform NHP viral infusions in an MRI scanner. We propose a CED modeling method which can assist any researcher in NHP neurosurgical CED planning.

Here we have developed a quantitative bench-side CED model which provides the users hands-on CED experience. Our bench-side model builds off of our recent qualitative infusion modeling work [19], as well as our *in vivo* NHP data [8], [9]. Bench-side CED models are usually comprised of dye infused into agar phantom, a clear gel with material properties similar to the brain [20], [21]. In this work we propose a similar model, but to our knowledge we are the first to base a bench-side model on *in vivo* MRI data of CED in NHP brains. We provide the MRI data and we present a calibration method for our model using the MRI data to ensure that reproduction of our quantitative method is practical for the field. Infusions into the primate brain are inherently coupled with surgical and experimental risks, however the toolkit presented here mitigates the risk factors of these procedures, such as cost, surgical time, and overall subject count, by providing easily accessible ways to plan CED experiments. We also utilize a radiolabel that can be co-infused with viruses that can be imaged in MRI 24 hours post-operatively, which empowers researchers without immediate access to an MRI scanner to identify issues and make any necessary surgical corrections in a timely fashion.

## 2. Materials and Methods

### 2.1 Subjects

Ten macaques were used for this study as described in Table 1. Ages ranged 5 – 11 years, and weights ranged 5.7 – 17.5 kg. Five females and four males were rhesus macaques (*Macaca mulatta*) and one male was a pigtail macaque (*Macaca nemestrina*).

**Table 1:** NHP and surgical data. NHPs are named for their infusion location(s). Medial temporal lobe and caudate nucleus (MTL). Hippocampus (HPC). Entorhinal cortex (EC), Tail of Caudate Nucleus (C).

| NHP Name | C1 | C2 | CT | T1 | T2 | MTL1 | MTL2 | MTL3 | MTL4 | MTL5 |
| --- | --- | --- | --- | --- | --- | --- | --- | --- | --- | --- |
| Infusion Target | Cortex |  | Cortex, Thalamus | Thalamus |  | MTL (HPC+C) |  | MTL (EC) | MTL (HPC+C) | MTL (HPC) |
| NHP Variety | Rhesus Macaque |  |  |  |  |  |  |  |  | Pigtail Macaque |
| Sex | M | M | M | F | F | F | F | F | M | M |
| Age (y) | 7 | 8 | 8 | 9 | 11 | 8 | 9 | 8 | 8 | 5 |
| Weight (kg) | 16.5 | 17.5 | 8.7 | 7.5 | 6.5 | 8.2 | 6.4 | 5.7 | 8.88 | 7.2 |
| Cannula Step<br>Tip Length (mm) | 1 |  | 1 (Cortex) and 3 (Thalamus) | 3 |  |  |  |  |  |  |
| Left Hemisphere Infusions (μl) | 50, 50, 50, 50 | 50, 50, 50, 50 | 50 (Cortex) | 140, 120 | 115, 85 | 15 | 20 | 10, 10 | 15 | N/A |
| Right Hemisphere Infusions (μl) | - | - | 246 (Thalamus) | 152, 111 | - | 15 | 20 | 10, 10 | 15 | 20 |
| Contrast Agent | Gadoteridol |  |  |  |  | Manganese |  |  |  |  |
| MRI Timing | Live | Live (2 infusions) | Live |  |  | Next-day |  |  |  |  |
| Institution | University of California, San Francisco |  |  |  |  | University of Washington, Seattle |  |  |  |  |
| NHP Alias | Monkey J [8] | Monkey G [8] | NHP-H [9], [23] | NHP-A [9] | NHP-B [9] | N/A |  |  |  |  |

### 2.2 Surgical procedures and MRI analysis

Three different neuroanatomical structures were targeted for CED infusions: cortex, thalamus, and the medial temporal lobe together with the caudate nucleus (MTL). Most of the cortical and thalamic infusions were validated with live MRI, while MTL infusions were validated with MRI the day following infusion. Table 1 contains surgical details for each subject.

Subsequent subsections describe surgical methods and MRI analysis related to cortical, thalamic, and MTL infusions. For all procedures, vital signs such as heart rate, respiration, and body temperature were monitored throughout.

#### 2.2.1 Cortical infusion

We have previously described the surgical methods for cortical CED procedures [8], [18] and briefly reiterate them here. The subjects were anesthetized with isoflurane anesthesia and mounted into an MR-compatible stereotactic frame. We made a sagittal incision on the left side of the head about 2 cm from the midline and peeled back the skin and musculature to reveal the skull. Then we used a trephine (25 mm diameter) to create a craniotomy over the sensorimotor cortex and performed a durotomy with ophthalmic scissors. A custom-made MR-compatible CED cannula guide was affixed to the skull with titanium skull screws or plastic screws and dental acrylic. The subjects were transported to the MR scanner and then positioned in the scanner (Siemens 3T). We filled the CED cannula guide with saline for MR visibility. A series of IV tubing was used to extend from a cannula positioned in the guide to a syringe pump (WPI UMP3, MICRO2T SMARTouch, SGE250TLL, Sarasota, FL, USA) located safely away from the MR-scanner. Our cannula was custom-made with a 1-mm stepped-tip similar to that described in section 2.3.1. We inserted the tip of the stepped-tip cannula about 2 mm below the surface of the brain in the sensorimotor cortex. The pump co-infused AAV-CamKIIa-C1V1-EYFP (Table 2; 2.5x10^12^ virus molecules/milliliter (vm/ml); UPenn vector core) and a gadolinium-based contrast agent (Table 2; ratio of 250:1, Gadoteridol, Prohance, Bracco Diagnostic Inc., Princeton, NJ, USA) into the brain at a starting rate of 1 µL/min, which was increased every few minutes up to 5 µL/min. After the majority of the volume had been delivered, the rate was reduced in the same stair-step method to end with a rate of 1 µL/min. After infusion, we left the cannula in place for 10 minutes and then removed the cannula. This infusion process was repeated multiple times. During the infusion process, multiple MRI images were taken to track the progress of the infusions. We used fast (2 minute) flash T1 weighted images (flip angle = 30°, repetition time/echo time = 3.05, matrix size = 128 x 128, slice thickness = 1 mm, 64 slices, Siemens 3T MR scanner). After the infusions and MR-scans were completed, we transported the subject back to the surgical suite where the cannula guide was explanted and the incision closed. In the cases of Monkeys C1 and C2, a titanium chamber and artificial dura were chronically implanted before the incision was closed. After recovery and optogenetic expression, Monkeys C1, C2, and CT were euthanized for immunohistochemical analysis of optogenetic expression [8], [22], [23].

**Table 2:**
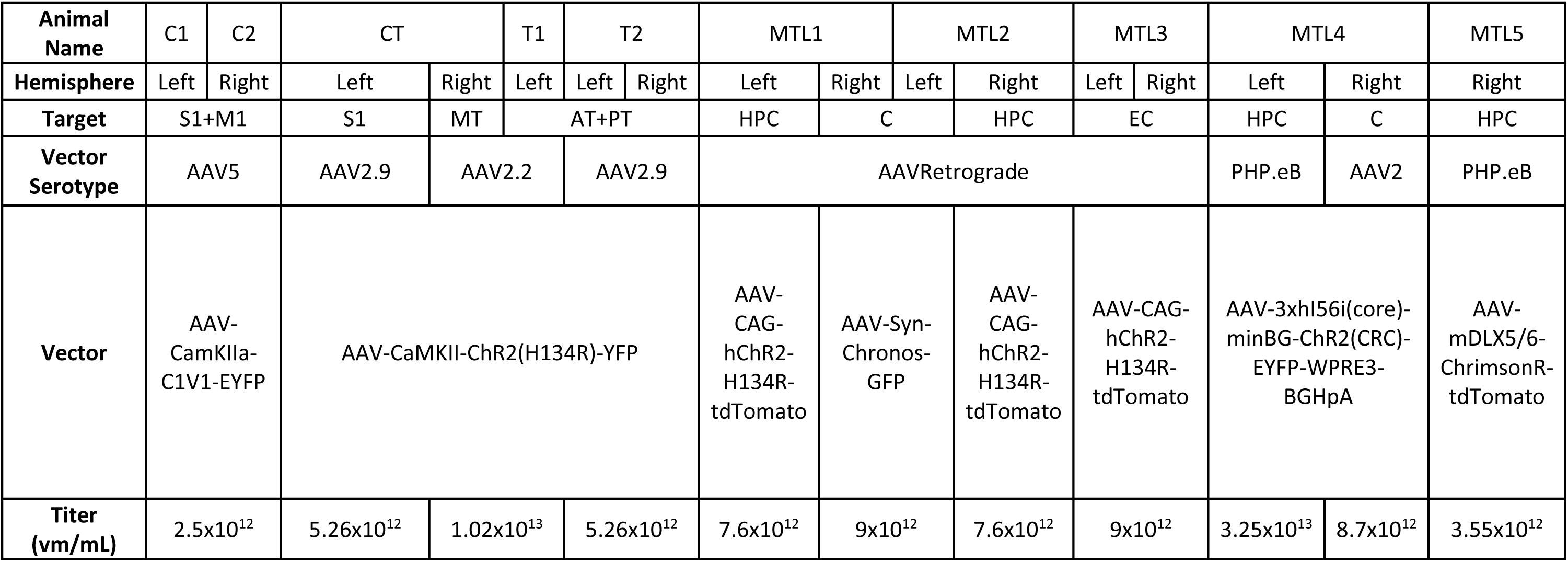
Virus Information. S1 (Primary Sensorimotor Cortex), M1 (Primary Motor Cortex), AT (Anterior Thalamus), MT (Medial Thalamus), PT (Posterior Thalamus), HPC (Hippocampus), C (Tail of Caudate Nucleus), EC (Entorhinal Cortex). vm/mL (virus molecules per milliliter).

#### 2.2.2 Thalamus infusion

We previously described the surgical details of the thalamic CED infusions [9], which are similar to the cortical infusion procedures. Briefly, the subjects were anesthetized and placed in a stereotax as in the cortical method, and we performed craniotomies (15 mm diameter). We implanted cannula guides and secured them with plastic screws and dental acrylic. We allowed the subjects to recover for two weeks before performing CED. After recovery, the subjects were anesthetized, placed in a stereotax and transported to an MR-scanner. We used the cannula guides to manually insert the 3-mm stepped-tip cannula (ClearPoint Neuro Inc. (formerly MRI Interventions), Solana Beach, CA, USA) to the targeted regions of the thalamus. We used a syringe pump (WPI UMP3, MICRO2T SMARTouch, SGE250TLL, Sarasota, FL, USA) to co-infuse multiple serotypes of AAV-CamKII-ChR2 (Table 2; Upenn vector core) with gadolinium-based contrast agent (Table 2; 2 mM Gadoteridol, Prohance, Bracco Diagnostic Inc., Princeton, NJ, USA) at rates between 0.5 µl/min and 3 µl/min while simultaneously performing fast (2 minute) flash T1 MR-scans (flip angle = 30°, repetition time/echo time = 3.05, matrix size = 128 x 128, slice thickness = 1 mm, 64 slices, Siemens 3T MR scanner). After all infusions were complete, the subjects were transported back to the surgical suite where the cannula guides were explanted and the incision closed. After recovery and optogenetic expression, Monkeys CT, T1, and T2 were euthanized for immunohistochemical analysis of optogenetic expression [9], [23].

#### 2.2.3 MTL infusion

To pilot the efficacy of expression in the medial temporal lobe (MTL), five viral vectors were infused into the brains of five subjects. Subjects MTL1-3 were infused with two retrograde viruses (gifted from Edward Boyden & Karel Svoboda; Addgene viral preps #59170-AAVrg & #29017-AAVrg, respectively) [24], [25]. Subjects MTL4&5 were infused with three GABA-selective viruses (Allen Institute for Brain Science & University of Washington, Seattle, WA.). See Table 2 for specific information about vectors and target regions for all animals.

The surgical details of the MTL infusions are similar to the thalamic infusion procedures. Briefly, after anesthesia and placing the subjects in the stereotax, burr holes were drilled and the 3-mm stepped-tip cannula (ClearPoint Neuro Inc. (formerly MRI Interventions), Solana Beach, CA, USA) was stereotactically lowered with a micro-manipulator arm through the burr hole to the targeted region of the MTL (hippocampus or entorhinal cortex), or the tail of the caudate nucleus. We used a syringe pump (WPI UMP3, MICRO2T SMARTouch, SGE250TLL, Sarasota, FL USA) to co-infuse optogenetic viral vector with manganese-based contrast agent (Mn^2+^, Millipore Sigma, Bulington, MA.) at rates between 1 µl/min and 5 µl/min. The final concentration of Mn^2+^ mixed with virus was 6.5mM and was specifically chosen as it is well under the limit for neuronal toxicity and interference with viral efficacy as identified by [26]. After all infusions were complete, the cannula was removed and the incision closed. In one case (MTL5), this procedure was performed using the Brainsight veterinary surgical robot (Rogue Research Inc., Montreal, QC, Canada). The day following infusion the subjects were again anesthetized and placed in an MR compatible stereotax, and 3-D MPRAGE sequences were taken in a 3T MRI Scanner to localize the manganese signal (Scanner: Phillips GE, Boston, MA., slice thickness: 0.35 × 0.35 × 0.5mm anisotropic & 0.5 isotropic voxels, repetition time/echo time = 2, Flip angle: 9 degrees). After eight weeks, animals were euthanized and perfused with 4% paraformaldehyde. Harvested tissue was then sliced into 50-μm sections and standard immunohistochemistry methods were employed to detect successfully infected target neurons of the various viruses with immunofluorescence.

#### 2.2.4 MRI volume extraction

The following infusion volume extraction procedure was performed on the MRI scans. For cortical and thalamic trials, the MRIs were taken throughout the infusion period. For MTL infusions, the MRIs were taken the day after infusion. We imported each MRI scan into MRI viewing software (Mango, Research Imaging Institute, UTHSCSA, San Antonio, TX, USA) and identified the location of the infusion – due to the contrast agent, this area had a higher contrast than the surrounding tissue. A spherical region of interest (ROI) was created and adjusted as necessary to encompass the infused volume. We shrink-wrapped the ROI in 3D with a threshold value below the intensity of the infusion location but above the intensity of the surrounding tissue. The threshold for the ROI was adjusted until it only contained the infusion volume and the final ROI was saved as a “.nii.gz” file. Next, we reloaded each ROI as its own image and generated an interpolated surface of the infusion volume. Finally, we measured the volume of the bolus using this interpolated surface in the MRI viewing software. In the case of live imaging during infusions, we mapped infusion trajectories by applying this protocol to successive MRI scans within an infusion trial.

### 2.3 Bench-side modeling

We developed a bench-side CED infusion technique using agar and custom-built cannulas. We also developed an image processing technique and statistical methods to analyze the agar data.

#### 2.3.1 Cannula production for agar infusions

We manufactured 1-mm and 3-mm stepped-tip cannulas (Supplemental Figure 1) with polyimide coated fused silica capillary tubing (Polymicro technologies, Phoenix, AZ, USA) for cortical and deep infusions respectively. These cannulas were created by sliding the smaller tubing into the larger tubing until the smaller tubing extended out of the larger tubing by 1 mm or 3 mm. This placement was then secured with cyanoacrylate (Super Glue Corporation, Liquid super glue, Ontario, CA, USA). For both cannulas, the inner tubing had an inner diameter of 320 µm and an outer diameter of 435 µm (part #1068150204) and the outer tubing had an inner diameter of 450 µm and an outer diameter of 673 µm (part #1068150625). The cannulas were the same as, or similar to, the cannulas used for our NHP CED experiments above.

#### 2.3.2 Agar phantom infusion

We prepared a solution of 1x phosphate buffered saline (PBS) and 0.6% powder mixture, where the powder mixture was comprised of agar powder (Benchmark Scientific, A1700, Sayreville, NJ, USA) and locust bean gum powder (Modernist Pantry LLC, Eliot, ME, USA) in a 4:1 ratio by mass respectively. We heated and mixed the solution in a microwave to dissolve the powder and poured it into molds to set. Setting occurred in a refrigerator for at least two hours. The molds were 3D printed with polylactic acid (PLA) filament and were designed to produce agar blocks with a 2 × 2 cm base and either 2 or 4 cm high. The agar phantoms were used shortly after setting or were refrigerated for up to one day for future use.

To prepare for CED infusion, we mounted a pump (WPI UMP3, MICRO2T SMARTouch, Sarasota, FL, USA) to a stereotactic arm (KOPF, 1460, Tujunga, CA, USA) attached to a stereotactic frame (KOPF, 1430, Tujunga, CA, USA). We filled the pump’s 250 mL syringe (WPI, SGE250TLL, Sarasota, FL, USA) with deionized (DI) water and attached it to the pump. The cannula was attached to the syringe with a catheter connector (B. Braun Medical Inc., part #332283, Bethlehem, PA, USA). All of the DI water was ejected from the syringe to fill the cannula with DI water. Undiluted yellow food coloring (McCormick yellow food coloring, Hunt Valley, MD, USA) was then drawn through the cannula and into the syringe. We positioned the agar phantom under the cannula so that it was centered. The agar phantom was oriented such that the side of the block which was not in contact with the mold during the molding process (i.e., the top side, which was the smoothest side of the phantom) was facing up and would be the side to receive the cannula insertion. The cannula was then lowered until the tip touched the surface of the agar phantom. We lowered the cannula manually with the stereotactic arm to a pre-specified depth, 2 mm deep for cortical infusions and 2 cm deep for thalamic and MTL infusions. We checked that the surface of the agar sealed around the cannula above the stepped-tip before proceeding with the infusion.

To image the infusion process, we positioned a digital single-lens reflex (DSLR) camera (Nikon D5300, Minato City, Tokyo, Japan) with a 35-mm lens (Nikon, AF-S NIKKOR 1:1.8G) and adjusted camera settings to clearly image the agar phantom edges, needle, and bolus. The ISO was set at 400, the shutter speed at 1/125, and the aperture at f/6.3. We used interval time shooting. We arranged a white backdrop to help with image processing and placed a ruler near the agar phantom and in-plane with the cannula for scale. We prepared a script in MATLAB (MathWorks Inc., Natick, MA, USA) to run the pump autonomously and in accordance with the infusion rates used in corresponding surgical infusions (Table 3). We started the script and the camera’s time interval shooting simultaneously. Representative cortical and thalamic infusion models are compared with MRI data as shown in Figures 1 and 2, respectively. Refinements to infusion techniques during preliminary trials included optimizing lighting conditions, and camera angle and placement with respect to the agar phantom.

**Figure 1.**
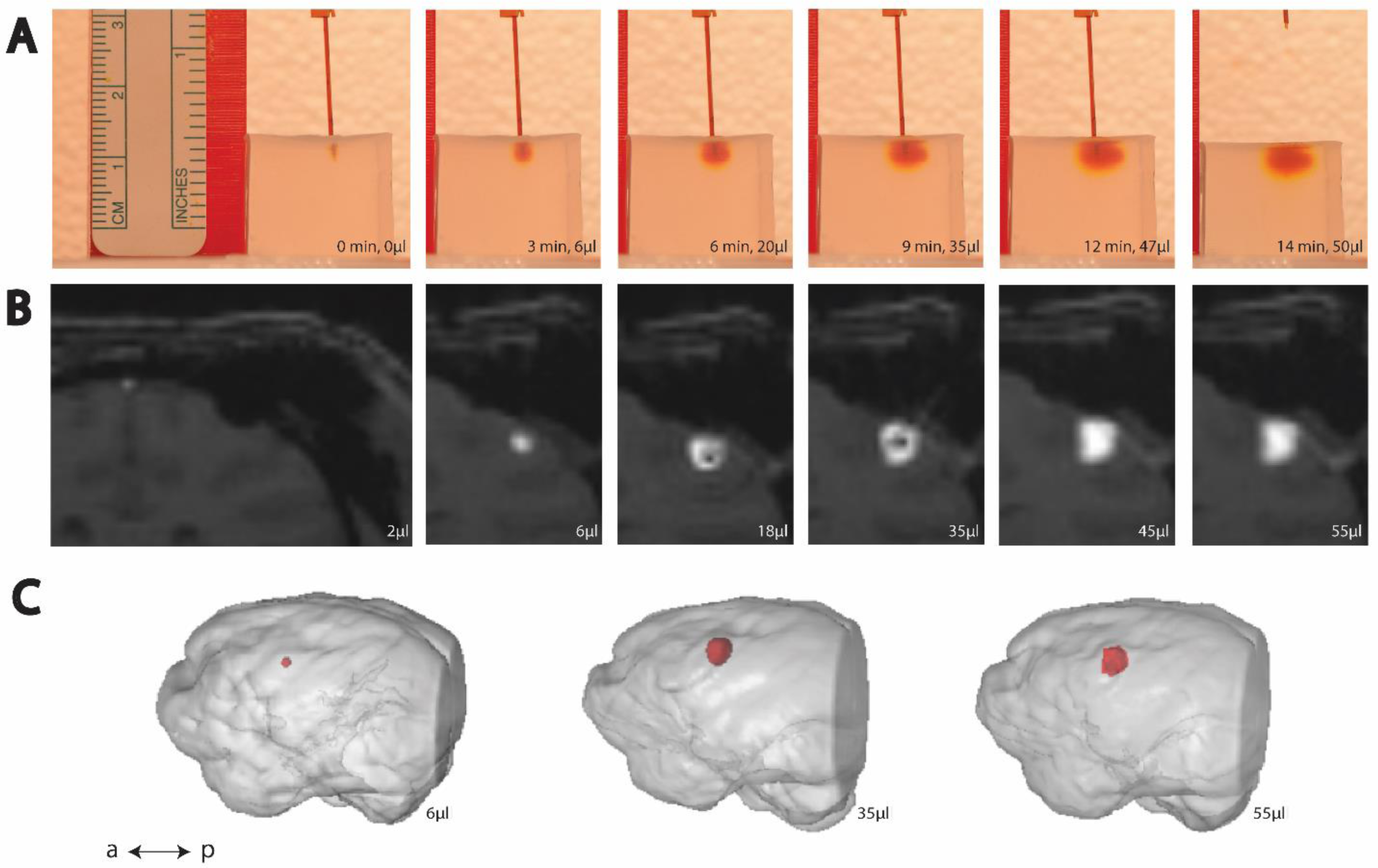
Time-lapse (left to right) of cortical CED. (A) Example trial of CED in agar phantom. (B) Example MRI visualization of CED in an NHP. (C) Post hoc reconstruction of NHP brain (grey) and infusion volume (red).

**Figure 2.**
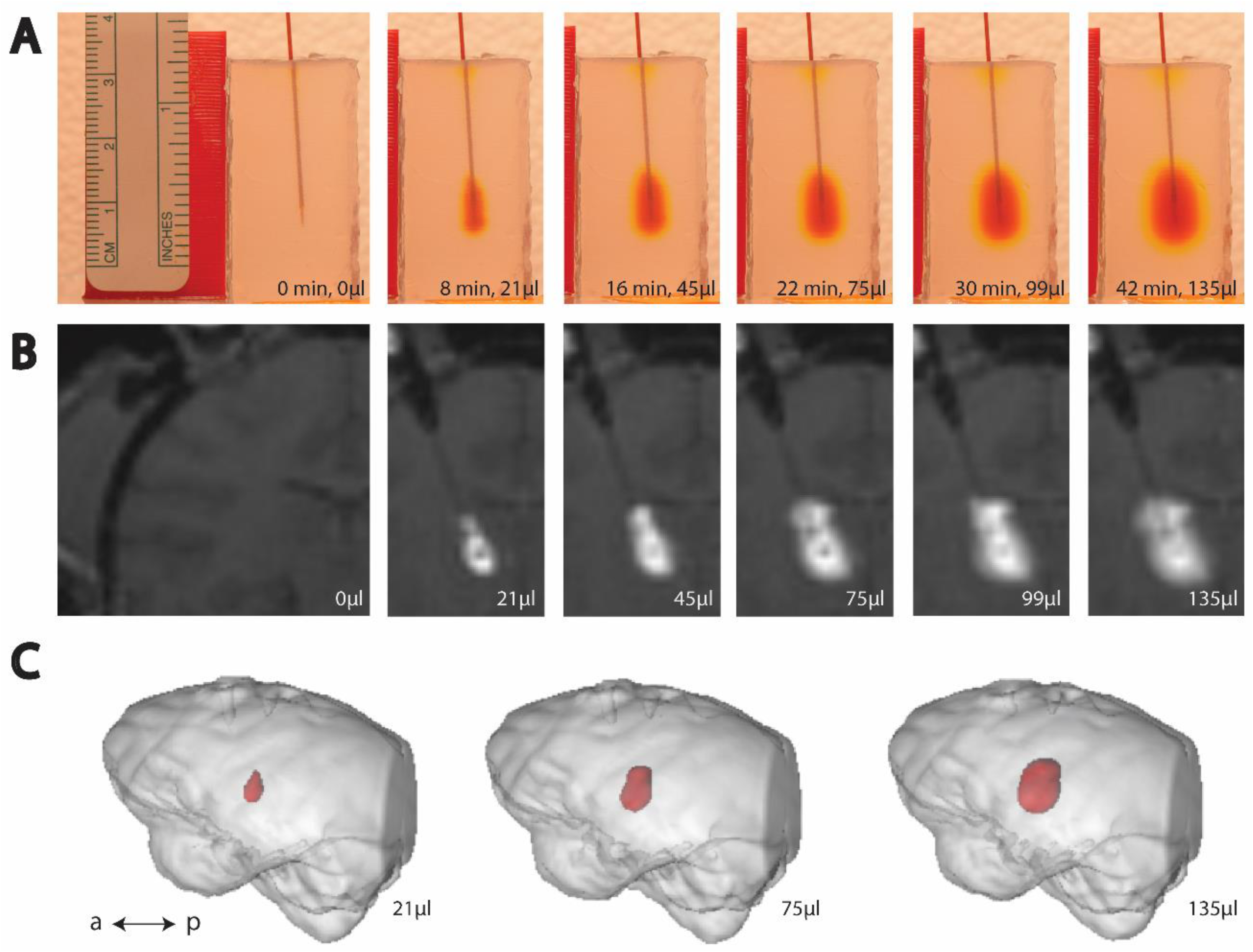
Time-lapse (left to right) of thalamic CED. (A) Example trial of CED in agar phantom. (B) Example MRI visualization of CED in an NHP. (C) Post hoc reconstruction of NHP brain (grey) and infusion volume (red).

**Table 3.** Gel infusion rates. Medial temporal lobe (MTL).

| Cortical<br>Infusion<br>Rates ($\mu\text{l}/\text{min}$) | Duration<br>(min) | | Thalamic<br>Infusion<br>Rates ($\mu\text{l}/\text{min}$) | Duration<br>(min) | | MTL Infusion<br>Rates ($\mu\text{l}/\text{min}$) | Duration<br>(min) |
| --- | --- | --- | --- | --- | --- | --- | --- |
| 1 | 1 |  | 1 | 1 |  | 1 | 1 |
| 2 | 1 |  | 2 | 1 |  | 2 | 1 |
| 3 | 1 |  | 3 | 80 |  | 3 | 3 |
| 4 | 1 |  | 2 | 1 |  | 2 | 1 |
| 5 | 6 |  | 1 | 1 |  | 1 | 1 |
| 4 | 1 |  |  |  |  |  |  |
| 3 | 1 |  |  |  |  |  |  |
| 2 | 1 |  |  |  |  |  |  |
| 1 | 1 |  |  |  |  |  |  |
| **Total infused:**<br>**50 $\mu\text{l}$** | **Total**<br>**time:**<br>**14 min** |  | **Total infused:**<br>**246 $\mu\text{l}$** | **Total**<br>**time:**<br>**84 min** |  | **Total infused:**<br>**15 $\mu\text{l}$** | **Total**<br>**time:**<br>**7 min** |

#### 2.3.3 Agar phantom image processing

We estimated volumes of distribution from photographs of the agar phantom infusion. A single color component was selected from the color images to reduce the three-dimensional image matrix into a two-dimensional image matrix. (Further description of the color component selection process is found below in section 2.3.4.) Then, we applied a threshold value to the remaining matrix to produce a mask, being a binary matrix of values indicating pixels above and below the threshold. (Specific methods for selecting this threshold value are found below in section 2.3.5.) We manually selected the cluster of binary values representative of the infusion bolus and erased the other clusters from the mask. In some cases, this was enough to isolate the bolus, but in other cases the mask appeared to depict the cannula together with the bolus. In these cases, the cannula was erased by masking all pixels above a manually selected point in the image such that the mask outlined the bolus alone. Representative images of the agar phantom image processing are shown in Figure 3. Finally, we converted the mask of the bolus to a volume by assuming an ellipsoidal form, where the ellipsoid was radially symmetric about the axis of the cannula. Similar to [27], [28], we took the height (*h*) and width (*w*) of the bolus and calculated the volume (*v*) of the associated ellipsoid with the equation

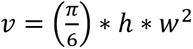

**Figure 3.**
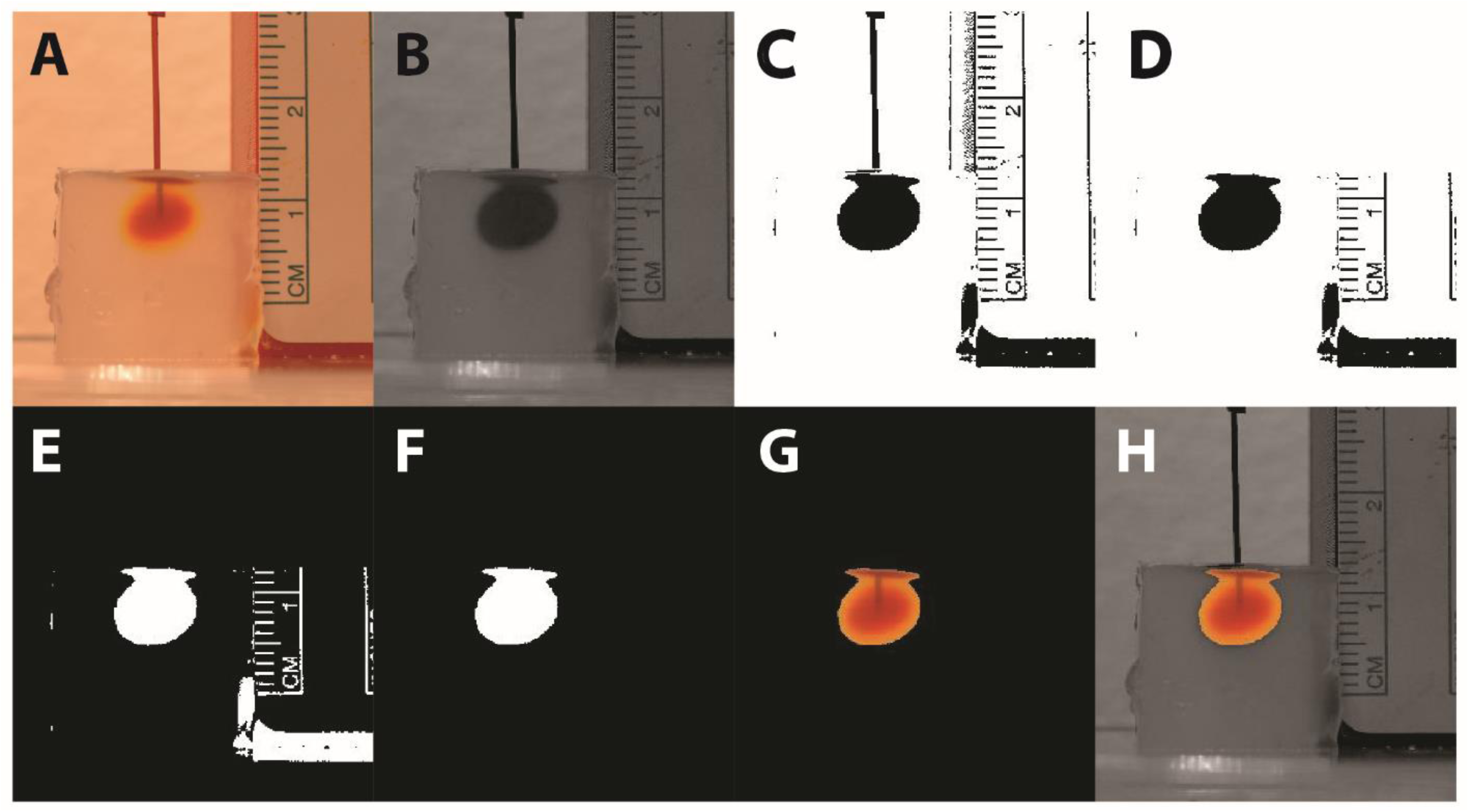
Image processing algorithm. (A) Original color image. (B) Single color component image. (C) Thresholded image. (D) Cannula erasure. (E) Binary inversion. (F) Deletion of all non-bolus pixels. This is the final image used for volume calculation. (G) Color image of bolus overlaid on final image for user reference. (H) Color image of bolus overlaid on the single color component image for user reference.

The volume estimation was converted to metric units based on the ruler in the original image.

We performed all agar image processing with MATLAB.

#### 2.3.4 Color component selection

Our color component selection is the process by which a color image with three color values per pixel (i.e., red, green, and blue) is converted to an image with one value per pixel. We tested the three different color components to determine which aligned the agar data most closely with the MRI data. All data of the same infusion type were processed with one selected color component. While image processing software often have functions built-in which will reduce images to a single value per pixel (e.g., the “rgb2gray” function in MATLAB), we found our color component selection process to be more effective.

#### 2.3.5 Threshold value selection

After the color image was converted to an image with one value per pixel (where values range from 0 to 255), we used a threshold value to distinguish which pixels had strong enough values to be included as part of the bolus. To select the threshold value, we identified a region of the image where heavy coloration faded to no coloration, then we selected a pixel from this region and use its value as the initial threshold. We performed the entire image processing procedure with this threshold value and plotted the agar data together with the MRI data to observe the quality of alignment. Based on the results, we selected a new threshold and repeated as necessary in an iterative fashion until the MRI data and the agar data were in alignment. Once aligned qualitatively, we compared the NHP volume data quantitatively with the agar volume data by using linear regression to determine the slope of each agar and NHP trial, as described in more detail in the next section (2.3.6). All data of the same infusion type were processed with one selected threshold value.

#### 2.3.6 Statistical methods

To compare the agar and NHP data for the cortical and thalamic trials, we required a quantitative method which would take into account correlations between data points within a given trial, and also compare the different groups of trials. Additionally, we required a method which would not assume a fixed slope between the volume infused by the syringe and the measured bolus volume. To this end, we used a linear mixed effects model with random slopes to fit each infusion trial, both in agar and in NHP data. All best-fit lines were restricted to passing through the origin (i.e., zero input volume and zero output volume). From this model, we calculated the interaction effect which is an approximation of the difference in average slope between the agar and NHP data. All linear mixed effect model calculations were performed with MATLAB’s built-in `“fitlme”` function.

All other statistical calculations were performed in MATLAB except the average and percent error values of Table 4, which were performed in Excel (Microsoft Corp., Redmond, WA, USA).

**Table 4.** Cortical and thalamic data.

| Cortical Infusion Slopes |  |  |  |  |
| --- | --- | --- | --- | --- |
| NHP | MRI Slopes ($\mu\text{l}/\mu\text{l}$) | | Trial | Gel Slopes ($\mu\text{l}/\mu\text{l}$) |
| C1 | 2.86 |  | 1 | 3.03 |
| C1 | 3.90 |  | 2 | 2.69 |
| C1 | 3.07 |  | 3 | 2.72 |
| C1 | 4.61 |  | 4 | 4.74 |
| C2 | 2.67 |  | 5 | 4.11 |
| C2 | 3.05 |  | 6 | 4.67 |
| CT | 3.07 |  | 7 | 4.23 |
|  |  |  | 8 | 3.10 |
|  |  |  | 9 | 4.81 |
|  |  |  | 10 | 3.82 |
| Thalamic Infusion Slopes |  |  |  |  |
| NHP | MRI Slopes ($\mu\text{l}/\mu\text{l}$) | | Trial | Gel Slopes ($\mu\text{l}/\mu\text{l}$) |
| T1 | 3.48 |  | 1 | 2.63 |
| T2 | 3.59 |  | 2 | 3.71 |
| T2 | 3.30 |  | 3 | 4.02 |
| CT | 6.38 |  | 4 | 2.66 |
|  |  |  | 5 | 3.17 |

## 3. Results

Not all agar infusion trials were successful, and outliers were omitted from statistical analyses. These outliers may have been produced by reasons such as dye leaking out through the catheter adapter or damaged agar phantoms.

### 3.1 Cortical and thalamic infusions

We iteratively selected color components and threshold values to align the agar data with our previously published cortical [8] and thalamic [9] NHP data (Figures 4 and 5, respectively). The green component and threshold value of 110 (43% of green component intensity range) were best for the thalamic infusions, and the blue component and threshold value of 67 (26% of blue component intensity range) were best for the cortical infusions.

**Figure 4.**
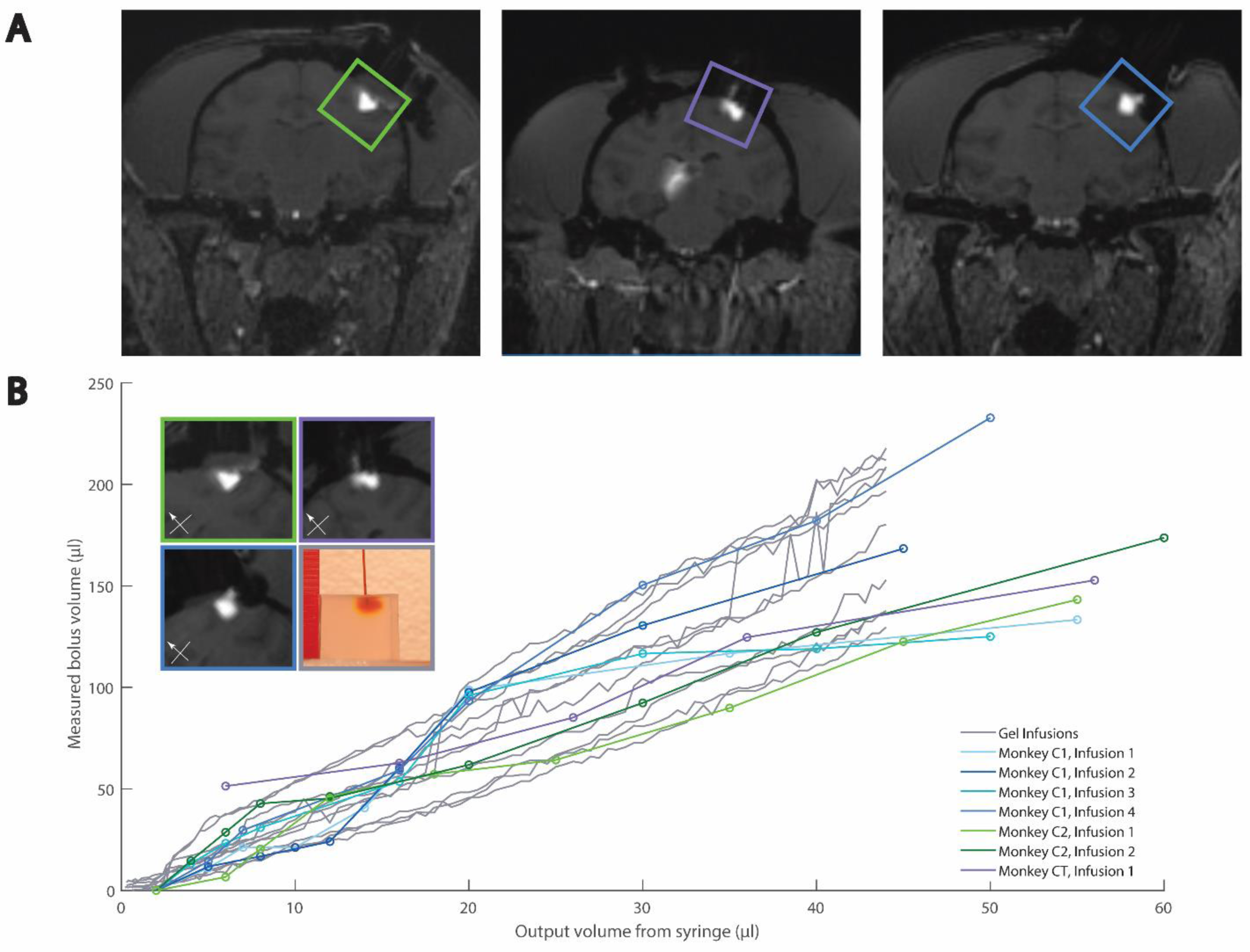
Comparison of agar and MRI cortical CED. (A) Example MRIs of cortical infusions. (B) Quantitative and example qualitative (inset) comparisons of agar and NHP data. Box colors in inset relate to images in (A) and traces in (B). MRI data has been previously published [8].

**Figure 5.**
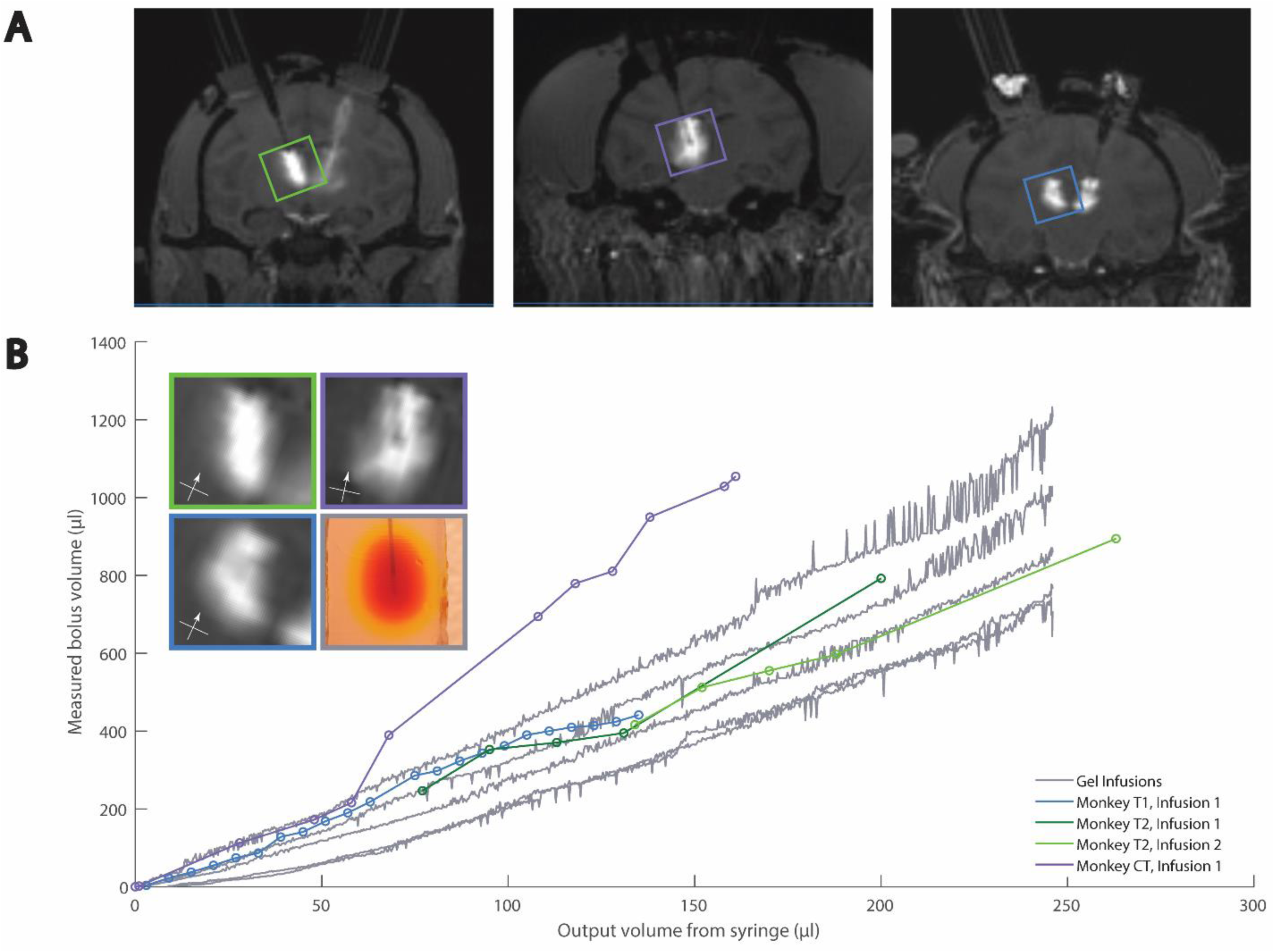
Comparison of agar and MRI thalamic CED. (A) Example MRIs of cortical infusions. (B) Quantitative and example qualitative (inset) comparisons of agar and NHP data. Box colors in inset relate to images in (A) and traces in (B). MRI data has been previously published [9].

When comparing the cortical agar data to the thalamic agar data, we observed that the agar and NHP best-fit slopes were similar (Figures 4 and 5). The cortical NHP data had slopes ranging from 2.9– 4.6 µl/µl, and the cortical agar data had a range of 2.7 – 4.8 µl/µl. Meanwhile, the thalamic NHP data had a range of 3.5 – 6.4 µl/µl and the thalamic agar data had a range of 2.6 – 4.4 µl/µl (Table 4). There is significant variation in the slopes of the best-fit lines of the agar infusion data, however, this variation reflects the variation in the NHP data (Figures 4B, 5B).

We observed that reduction of flow rate at the end of the cortical agar trials led to greater increase in bolus volume with respect to infused volume, i.e., the plots steepen near the end of the infusion protocol (Supplemental Figure 2). This is not characteristic of NHP cortical data, so we omitted the agar cortical data from 44 µl to 50 µl from statistical analysis and Figure 4 because it was not characteristic of NHP cortical data. Further description may be found in the discussion (section 4.4).

For the cortical and thalamic trials, we used a linear mixed effects model with random slopes to fit each infusion trial, both in agar and in NHPs. This model produces an interaction effect of 1.1 and -0.5 for cortical and thalamic infusions respectively. These values are close to zero in comparison with the aforementioned ranges of the slopes, and these values have magnitudes less than the slope ranges, indicating that the agar and NHP data differ only mildly and that our agar phantom is a good representation of the NHP data for cortical and thalamic infusions.

### 3.2 MTL infusions

Our MTL NHP data were highly variable, and upon analysis of the MRI scans, we observed that four of the nine infusions displayed a bolus in the MTL as expected (Figure 6A). We only modeled these four successful MTL NHP infusions and omitted the remaining five infusions, which are discussed below (section 3.2.1).

**Figure 6.**
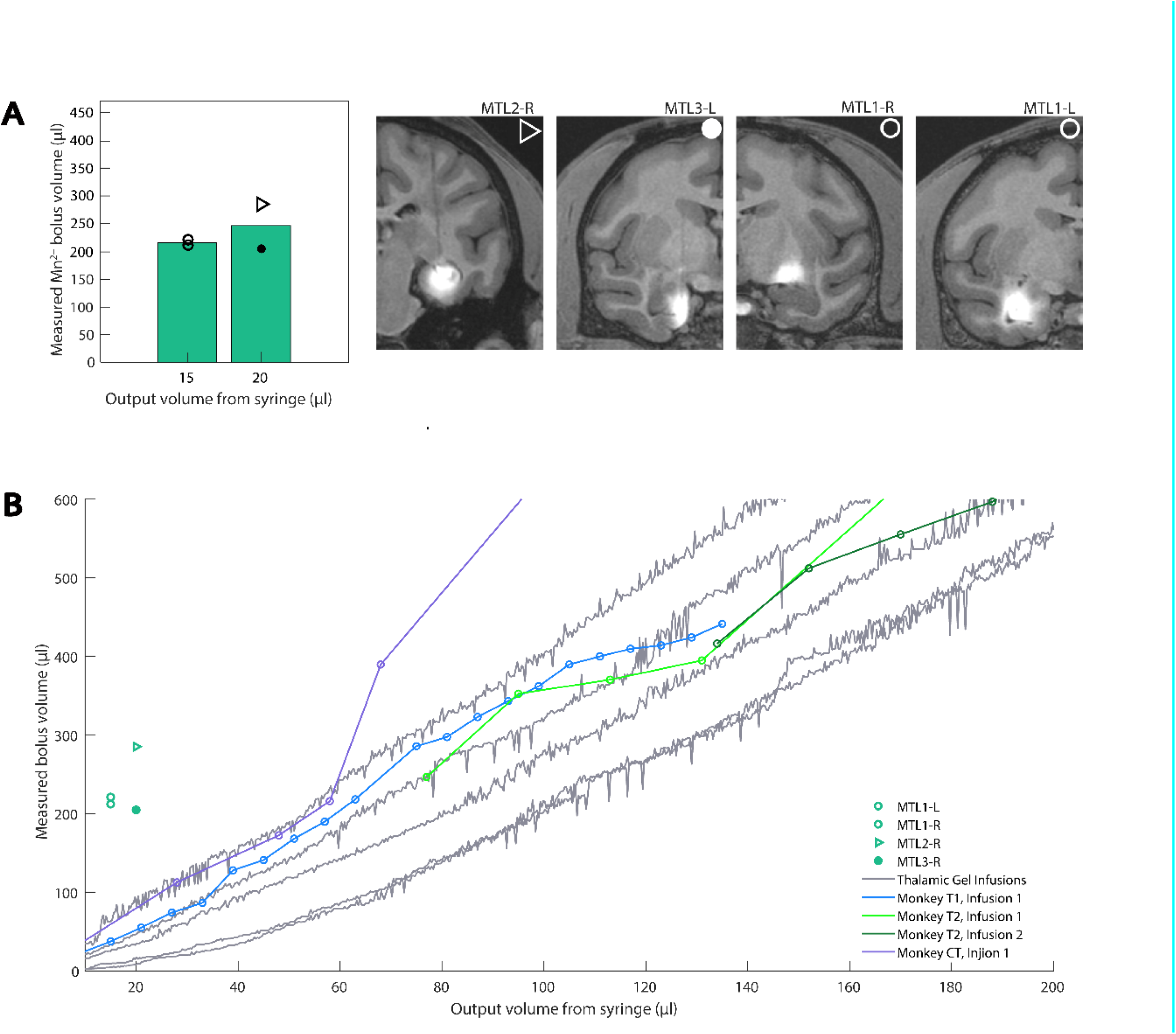
MTL MRI data compared with both agar and MRI thalamic data. (A) Deep CED infusions were made into the hippocampus, entorhinal cortex, and the tail of the caudate nucleus. Bar plots show the mean value of measured Mn^2+^ bolus seen in next-day, post-operative MRIs, with individual data points overlaid. Corresponding MRI slices in the coronal plane are shown for infusions into deep brain areas which are visually similar to cortical and thalamic data. Shape labels correspond to each subject contributing to the data. (B) MTL MRI data compared with both agar and MRI thalamic data. Thalamic MRI data has been previously published [9].

Because the thalamic and MTL NHP infusions used similar cannulas and differed chiefly in the infused volume, we initially compared thalamic agar bolus volumes with MTL NHP bolus volumes (Figure 6B). Counter to our expectations, the MTL NHP data did not align with either the thalamic NHP or agar data. We recognized that while the cortical and thalamic MRI scans were collected during CED infusion, the MTL MRI scans were collected the next day. Therefore, we reasoned that a CED-generated bolus may diffuse overnight and thus be displayed as a larger bolus in the NHP when MRI is performed the day after infusion. With this in mind, we proposed that diffusion of food coloring in agar after a standard CED protocol would approximate our MTL data. We collected data following the end of 15-µl infusions into agar and observed that the agar data closely modeled the NHP data after approximately 29 minutes of diffusion following the completion of the infusion (Figure 7). We calculated the mean of the NHP infusion volumes and the mean of agar infusion volumes selected from approximately 29 minutes after infusion completion (Table 5), and report a 3.5% percent error between the two data sets. Given the biological context of our model, this error is small enough to safely conclude that 29 minutes of diffusion in our agar phantom, following CED, approximates next-day MRI results of NHP MTL infusions.

**Figure 7.**
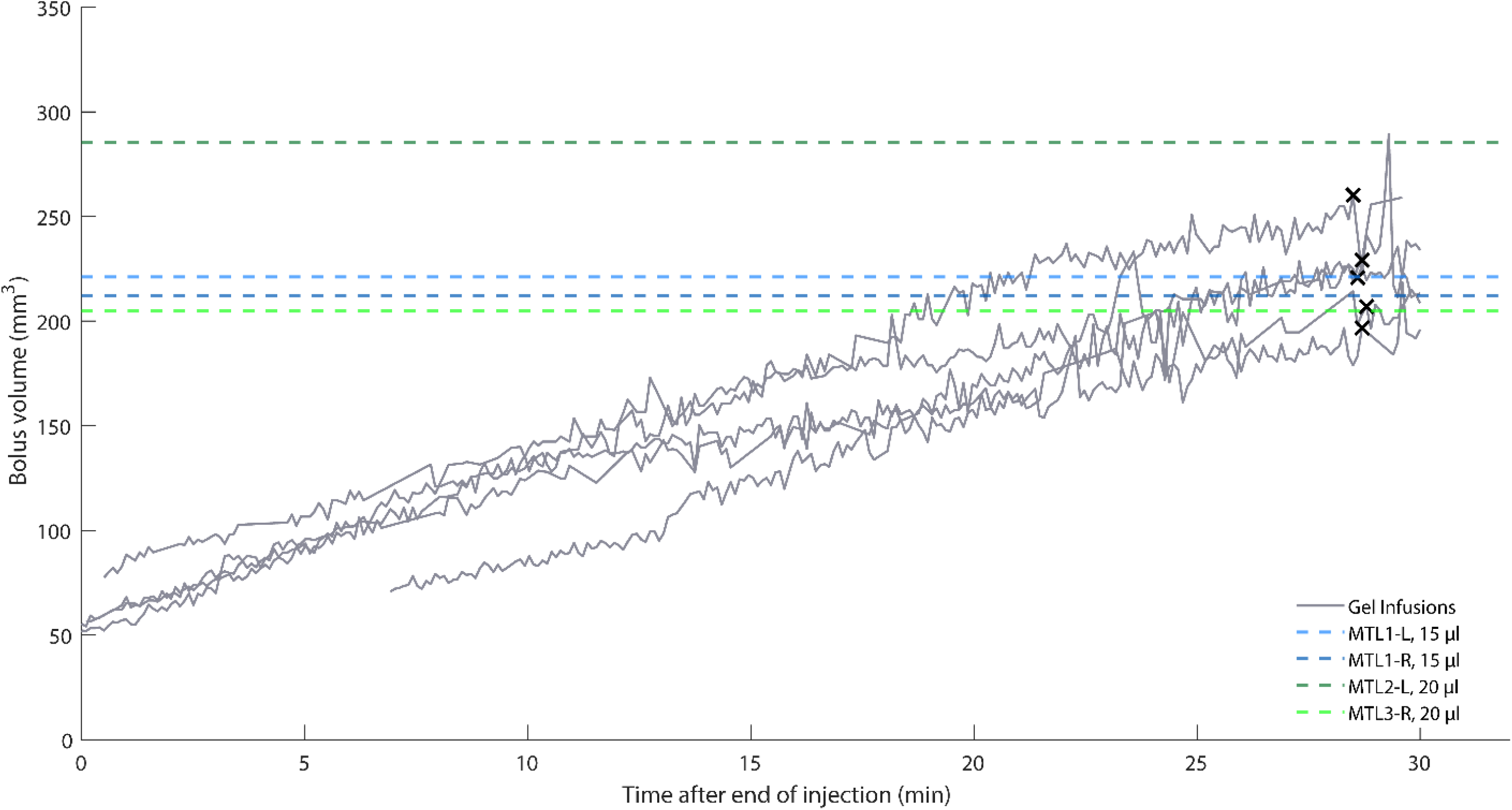
MTL MRI data compared with MTL agar data after CED completion. Some trials lack data early in the diffusion process due to the bolus being difficult to identify. Black X’s indicate the final point used for statistical analysis from each agar trial. The final chosen points may be shifted slightly to avoid noise spikes. All agar infusions were 15 µl.

**Table 5.** MTL data.

| **NHP,**<br>**Hemisphere** | **Infusion**<br>**Volume (μl)** | **Measured Volume**<br>**(μl)** |
| --- | --- | --- |
| MTL1, left | 15 | 221.1 |
| MTL1, right | 15 | 212.2 |
| MTL2, left | 20 | 285.3 |
| MTL3, right | 20 | 204.7 |
| **Average:** |  | **230.8** |
| **Gel Trial** | **Infusion**<br>**Volume (μl)** | **Measured Volume**<br>**(μl)** |
| 1 | 15 | 196.7 |
| 2 | 15 | 229.2 |
| 3 | 15 | 260.2 |
| 4 | 15 | 220.7 |
| 5 | 15 | 207.0 |
| **Average:** |  | **222.8** |
| **% Error:** |  | **3.5** |

The green component and threshold value of 100 (39% of green component intensity range) were best to model the MTL infusions. We used a t-test to compare the two NHP data points for 15-µl infusions and two data points for 20-µl infusions, and because the two populations were not statistically significant (p = 0.56), we pooled the four NHP data points for comparison with agar data. Significant noise was observed in the MTL agar data, more so than in cortical and thalamic agar infusions. The additional noise was possibly due to smaller infusion volumes. We addressed the noise by fitting a line with linear regression to each agar infusion trial and removing data points greater than 1.1 times the corresponding value on the best-fit line.

#### 3.2.1 MTL data omitted from agar modeling

Five of the nine NHP MTL infusions were excluded from the analysis since the next day MRI did not confirm infusion into the MTL regions (Figure 6, Supplemental Figure 3). These failed infusions mostly likely occurred due to the complex shapes of the hippocampus and neighboring structures, which contrasts with the structures of the cortex and thalamus. In three of the five unsuccessful MTL cases, the boluses were very small, which suggested either puncturing into the hippocampal fissure, or infusing deep in the dentate gyrus, depending on the injection depth (Supplemental Figure 3).

Penetration of the fissure resulted in contrast agent and virus partially escaping our injection target, indicative of unsuccessful CED. By contrast, deep dentate gyrus infusions were limited to a small portion of the hippocampus, limiting the amount of diffusion observed when compared to our gel model. In the remaining unsuccessful case, the bolus was very large and extended well outside of the volume of the hippocampus, indicating that contrast agent refluxed along the track of the cannula (Supplemental Figure 3). We have excluded these five unsuccessful MTL CED data points from our models, but these negative results are presented to highlight the value of *in vivo* verification of deep injection surgeries.

### 3.3 Histological analysis

For cortical and thalamic infusions, spread of the MR contrast agent modeled the volume of expression of the optogenetic viral vector, as previously analyzed and reported [8], [9]. We used different constructs, including retrograde viruses, for our MTL infusions. Because of the variability of the resulting expression, further experiments using a single virus known to express well in these regions is necessary to confirm whether next-day, Mn^2+^ MRI signal mirrors expression. Nevertheless, the Mn^2+^ MRI confirmed our targeting in vivo. Additionally, preliminary evidence from the successful local infection in case MTL4-L suggests a close match between Mn^2+^ signal and immunofluorescence (Figure 8).

**Figure 8.**
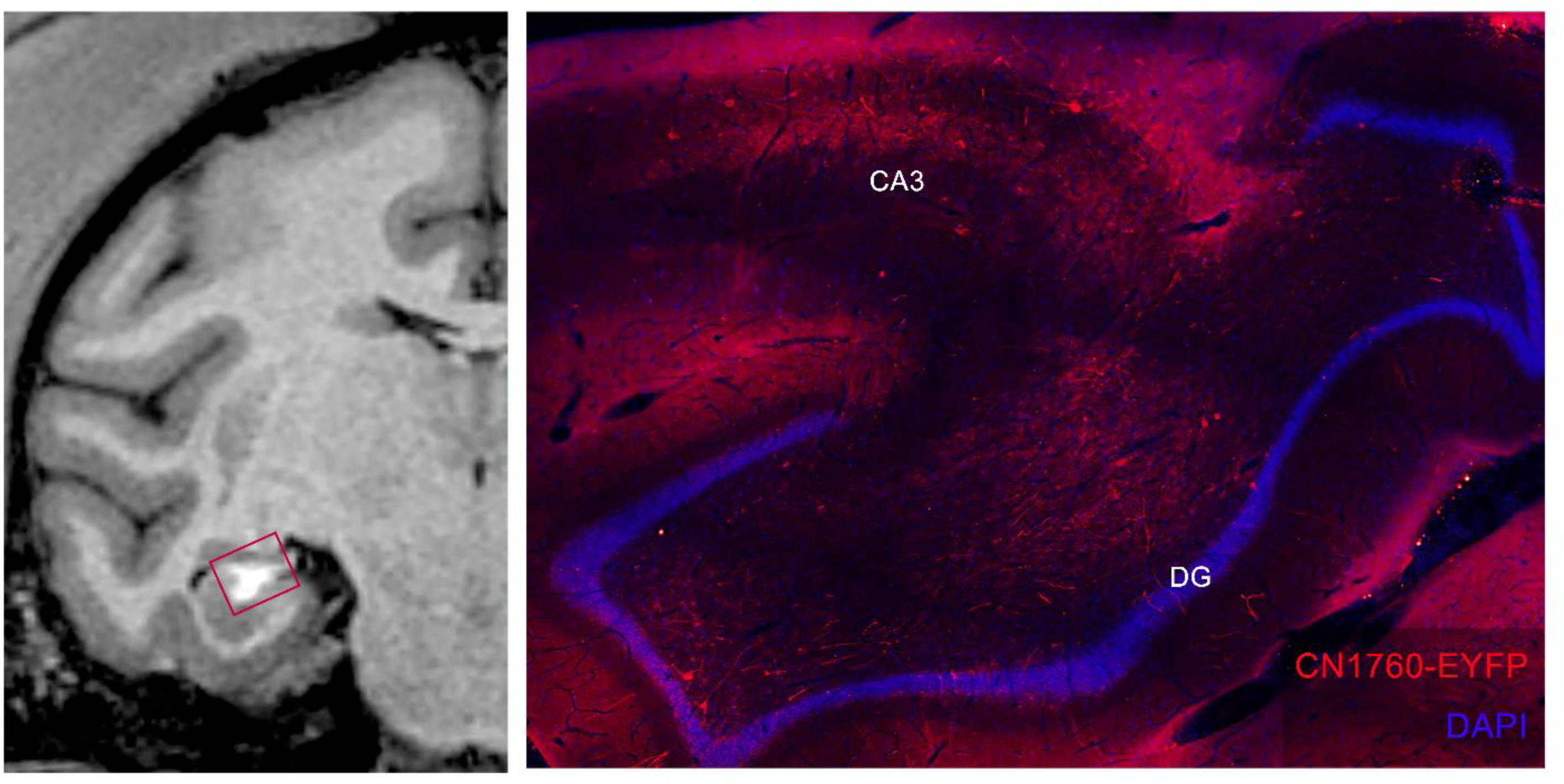
Comparison of Mn^2+^ MRI signal (left) and local expression of virus (right) after a hippocampal injection in case MTL4-L. Right-imaged region highlighted with red square in MRI (left). Our virus used a GABA-specific enhancer to selectively target interneurons, as evidenced by a lack of red pyramidal cell body labeling in CA3 and granule cell body labeling in the dentate gyrus (DG). We tagged infected interneurons and associated fibers with red-shifted fluorophores (anti-EYFP, ThermoFisher, Waltham, MA., USA) and cell bodies were non-selectively labeled using a DAPI stain (ThermoFisher, Waltham, MA., USA). (Viral construct CN1760: trAAV-3xhI56i(core)-minBG-ChR2(CRC)-EYFP-WPRE3-BGHpA (Paul Allen Institute, Seattle, WA., USA).)

## 4. Discussion

Targeted neural manipulations, such as those achieved via optogenetics, are revolutionary techniques for investigating circuit-level communication in the brain and have the potential to influence novel neurotherapeutic technologies in humans. NHPs are the keystone model for validating these techniques because of their similarity to humans, but they are a scarce resource that does not allow for experiments with many unknowns. As such, the risks associated with the viral infusions necessary for most genetic manipulations have led to a lack of uniformity in experimental design and much trepidation in engaging this type of research [29]. In this work, we developed a simple and efficient pipeline to ameliorate a number of these concerns. Our method improves upon past models [7], [20], [21] by quantitatively matching agar data with *in vivo* infusions of viral particles co-infused with MRI contrast agents, which serve as a proxy of the effective infusion volume. Further, we observe that certain contrast agents can signal the location of infusions up to 24 hours post-infusion, which is a greater time delay for bolus localization than previously demonstrated [26]. Taken together, the methods presented here serve as an accessible and inexpensive protocol to plan the optimized spread of infusions bench-side and validate the spread and accuracy *in vivo*, significantly reducing the number of unknowns that hinder confidence during circuit-manipulation experiments.

Agar has been previously established as a model of intraparenchymal neural tissue and is now commonly used as a medium for simulating infusion procedures [7], [20], [21]. Our work builds on prior studies by presenting a data-driven method developed from *in-vivo* results. To our knowledge, the model proposed here is the first to provide a quantitative method of fitting agar infusion data with NHP CED data collected with live MRI. Because of this, our model serves as a more accurate guide for selecting infusion parameters for future *in vivo* infusions targeting a wide array of brain areas when compared to other simulations.

Our work lends itself well to case-by-case methodological refinements: for example, researchers may consider alternative cannulas, infusion protocols, etc. to cater to their goals. Additionally, our method is designed to facilitate replication by other labs with its simplicity in both materials and methods. We also recognize that labs replicating our work are unlikely to implement agar infusion imaging setups identical to our own. With this in mind, we designed our image processing technique to be easily adaptable to different imaging setups. We provide NHP MRI bolus volume data (see Supplementary Materials) to which other labs may align their own agar infusion data, using our presented method. We have also made our custom code freely available (see Data Availability Statement), and the code is written in MATLAB which is widely used by researchers and is straightforward to adapt for applications akin to ours.

### 4.1 Diversity of Modeled Structures

To maximize flexibility in accurately predicting spread of CED infusions in a variety of brain areas, we used *in-vivo* data from cortical, thalamic, and MTL infusions of viruses co-infused with MRI contrast agent. Our cortical and thalamic procedures, which represented shallow and deep infusions into large brain structures, utilized live MRI taken during surgery. In addition to qualitative agreement between these infusion types, the ranges of MRI data best-fit slopes were similar (2.9 – 4.6 µl/µl for cortical and 3.5 – 6.4 µl/µl for thalamic) which suggested that the agar models for cortical and thalamic infusions may be similar as well. This proved to be the case: Despite differences in cannula design, depth of insertion, and infusion protocol, the models for both cortical and thalamic CED were generated with our same presented method and aligned well to the *in vivo* data (agar model best-fit slopes were 2.7 – 4.8 µl/µl for cortical and 2.6 – 4.4 µl/µl for thalamic). The cortical and thalamic models did differ in the parameters used during image processing (cortical: blue component, threshold = 67; thalamic: green component, threshold = 110), but this demonstrates that our method is robust to variations in agar infusion processes. Our successful MTL cases differed from the previous two conditions and represented infusions into more limited and more difficult to access deep structures. Additionally, they differed in their post-operative MRIs in that scans were taken ∼20 hours after infusion. We found that our initial hypothesis of these data aligning to the thalamic cases was invalidated. However, after accounting for diffusion expected from the delay in imaging, our model successfully aligned to all three conditions.

### 4.2 Insights from MTL Infusions

Because of its unfurled shape in primates compared to rodents [30], standard injections into the NHP hippocampus are laborious and often require either multiple craniotomies and penetrations [31] or penetrating through the long-axis of the structure and periodically injecting while retracting [32] to try and maximize coverage. To inject into this structure more efficiently, we leveraged the unique ability of CED to deliver a large bolus with a single infusion in our MTL CED group, which to our knowledge is the first set of CED infusions delivered to this area in NHPs. Because of the exploratory nature of these experiments, we experienced challenges that made delivery into this structure and subsequent imaging of our contrast agents more difficult. Infusion MTL2-L was a case of mass reflux due to an error made during the infusion delivery. MTL3-R and MTL4-R represent issues in targeting. Because much of the hippocampus is separated from the rest of the brain by ventricular spaces except laterally, targets made too shallow or too deep will leak into those spaces and either dissipate away or reflux upwards. For similar reasons, posterior-medial injections—for example targeting the intermediate dentate gyrus— produced more isolated boluses (cases MTL4-L, MTL5-R). Our successful cases were qualitatively similar to our cortical and thalamic data because they were delivered to larger, more anterior regions in the genu of the hippocampus, or to large neighboring regions such as the entorhinal cortex or tail of the caudate nucleus. For areas such as MTL where targeting needs to be very precise, we strongly recommend the use of MRI validation of injection either during infusion or the next day.

### 4.3 MRI scan parameters for successful contrast label visualization

Our novel MTL infusions were also the first to utilize co-infused manganese to localize viral infusions in MRI scans acquired ∼20 hours post-operatively. Despite the differences in employed contrast agent, scan acquisition timing, and even the scanner used between this group and our cortical and thalamic infusions, we observed a few similar parameters for successful contrast imaging in all MRI scans for all groups. Specifically, all scans were T1-weighted scans with a repetition time/echo time ratio around 2 to 3 and flip angles from 9 to 30 degrees. Analysis of other studies employing similar manganese-enhanced MRI protocols either to image viral injections delivered at shorter delays [Fredericks cite] and at much longer delays for *in-vivo* tract tracing [33] also closely mirrored the majority of parameters used in these experiments, suggesting a range of optimized parameters for imaging T1-weighted MRI contrast agents.

### 4.4 Diffusion versus convection using agar

It is important to note that while agar is a good model of CED, agar’s rate of diffusion differs from the rate of diffusion in the brain. This factor became apparent when we observed that reduction of flow rate at the end of the cortical agar trials led to greater increase in bolus volume with respect to infused volume, i.e., the plots steepen near the end of the infusion protocol (Supplemental Figure 2). This is consistent with our observation that diffusion will continue to cause the bolus to grow in agar after the end of our infusion trials. With this in mind, we concluded that diffusion and convection both contribute to bolus size in agar to varying degrees during CED. However, we propose the relative contributions of diffusion and convection were skewed when the flow rate was reduced at the end of the protocol, thus allowing diffusion to contribute more heavily to the bolus size in agar. To prevent the best-fit lines of the cortical agar data from being skewed due to this effect, we omitted the data from 44 µl to 50 µl from statistical analysis and Figure 4. This effect was not observed in the thalamic agar infusions likely due to the lower maximum rate of infusion (3 µl for the thalamic agar protocol, as contrasted with 5 µl for the cortical agar protocol) in conjunction with the short duration of infusion at the lower rates and small amount of volume infused during the flow rate reduction at the end of the protocol. This effect was not observed *in vivo* cortical or thalamic injections collected with live MRI. We observed diffusion in our *in vivo* MTL data, but the data were collected with our next-day imaging technique, thus allowing sufficient time for diffusion. We show that MTL next-day imaging data can be modeled with our agar phantom when we allow 29 minutes of diffusion following the end of the infusion protocol. This highlights that the speed of diffusion in agar differs from that of the brain. Also, this is in agreement with our observation that cortical and thalamic NHP data collected with live MRI did not exhibit high levels of diffusion at the end of the protocols, in contrast with cortical agar data.

### 4.5 Technical Considerations

We encountered some issues during agar infusions. The most common issues faced when refining infusion techniques were damaging the agar during its extraction from the mold, such that no smooth surface was available for imaging, and reflux of the dye during the infusion. While agar preparation became more efficient with practice, we suggest that custom, flexible silicone molds be considered in lieu of our 3D printed molds to more easily produce undamaged agar phantoms. Reflux issues can arise in agar, neural tissue, and other media and are less easily mitigated because the cause of reflux is not always obvious. In some cases, reflux may be related to the quality of the seal between the media and cannula, which is difficult to assess visually even in transparent media such as agar. Additionally, we hypothesize in some cases that the cannula gets clogged with the media during insertion. To address this potential issue, we suggest a low flow rate during insertion to avoid clogging.

As previously mentioned, the agar data are aligned to *in vivo* MRI data with an iterative process of parameter selection. The iterative alignment process allows researchers to fine-tune their image processing parameters to overcome potential differences in lighting, camera placement, etc. While our agar image processing techniques are effective, we acknowledge that software refinements may be attained. Our process is currently semi-automated yet we expect it could be more fully automated in future work. Improvement opportunities may also exist in the refinement of our volume estimation formula, and the characterization of the bolus shape.

### 4.6 Ethical Considerations

Despite the limitations presented, our methods will allow for the development of more efficient and effective CED procedures. We can inexpensively plan the expected spread of our infusions with our data-driven model and validate our surgical targeting rapidly without the need to perform infusions in an MRI scanner. Critically, our method is not only quantitative and data-based, but also designed to aid surgical planning with its visual, hands-on nature. Our bench-side modeling technique serves to increase the likelihood of success in NHP CED experiments, thus refining animal research processes and reducing the number of animals required for experimentation, both of which are key ethical considerations in animal research and included in the 3Rs [cite]. Our model was capable of simulating both cortical and deep infusions (limitations discussed above). Because of this, our method provides a generalized surgical preparation technique to all researchers regardless of region of interest, and particularly to research groups which do not have the facilities or resources required to perform live MRI during CED infusions. Our novel next-day MRI data additionally serve to showcase a post hoc infusion confirmation method that improves upon previous work [26] to highlight verification of infusion placement ∼20-hours post-operatively. This method supplements our proposed modeling technique and is a welcomed alternative to live-MRI, which requires specialized equipment and facilities often unavailable to researchers. In sum, we propose our method as an additional way of applying the principles of replacement, reduction, and refinement (3Rs) to injections in NHPs [34]. We recommend NHP CED infusions be modeled in advance of surgery with our proposed method to reduce the number of animals, replace an excess of pilot procedures with artificial simulations, and refine the overall technique to reduce harm. We also suggest the results be confirmed after surgery with MRI if live-MRI is not feasible during infusion. Finally, our work is designed to be highly flexible. While our methods are specifically prepared for NHP experiments involving optogenetic actuators, we expect that our method would also be generally effective for modeling CED of optogenetic sensors, pharmaceutical compounds, and other therapeutic agents in large brains.

## Supplementary Materials

We provide three supplementary figures and a CSV file of the MRI data points.

## Acknowledgements

We would like to thank all of the staff from University of Washington, Seattle, and University of California, San Francisco who provided animal care and surgical support. We thank Sabes lab for use of their previously published data. We thank Spencer Hansen, Serge Aleshin-Guendel, and Tianyu Zhang for their help with statistical analyses. We thank the Viral Technology team at the Allen Institute for Brain science, Ximena Opitz Araya, Shane Gibson, and Greg Horwitz for assistance with virus production. Finally, we thank Toni Haun, Karam Khateeb, and Megan L. Jutras for their technical help.

## Author Contributions

Conceptualization, E.A.B. and A.Y..; Methodology, D.J.G., A.D.G., W.Y.A., W.K.S.O., A.G.J., E.A.B., and A.Y..; Software, D.J.G., A.D.G., W.Y.A., and W.K.S.O.; Validation, D.J.G. and A.D.G.; Formal Analysis, D.J.G., W.Y.A., and W.K.S.O., A.Y.; Investigation, D.J.G., A.D.G., W.Y.A., W.K.S.O., A.G.J., E.A.B., and A.Y.-S.; Resources, E.A.B., A.Y., and J.T.T..; Data Curation, D.J.G., A.D.G., W.Y.A., W.K.S.O., A.Y. and A.G.J.; Writing – Original Draft Preparation, D.J.G., A.D.G., W.Y.A., and W.K.S.O.; Writing – Review & Editing, D.J.G., A.D.G., W.Y.A., W.K.S.O., A.G.J., E.A.B., and A.Y.; Visualization, D.J.G., A.D.G., W.Y.A., E.A.B., and A.Y.; Supervision, E.A.B. and A.Y.; Project Administration, E.A.B. and A.Y.; Funding Acquisition, E.A.B. and A.Y.

## Institutional Review Board Statement

All animal care and experiments performed at the University of Washington were approved by the University of Washington’s Office of Animal Welfare, the Institutional Animal Care and Use Committee, and the Washington National Primate Research Center (WaNPRC). All animal care and experiments performed at the University of California, San Francisco were performed under the approval of the University of California, San Francisco Institutional Animal Care and Use Committee and were compliant with the Guide for the Care and Use of Laboratory Animals.

## Informed Consent Statement

Not applicable.

## Data Availability Statement

MRI files, agar data, and histological data will be made available upon reasonable request. MATLAB code has been made publicly available: https://bitbucket.org/yazdanlab/ced_protocol_and_analysis/src/master/

---

**Supplemental Figure 1.**
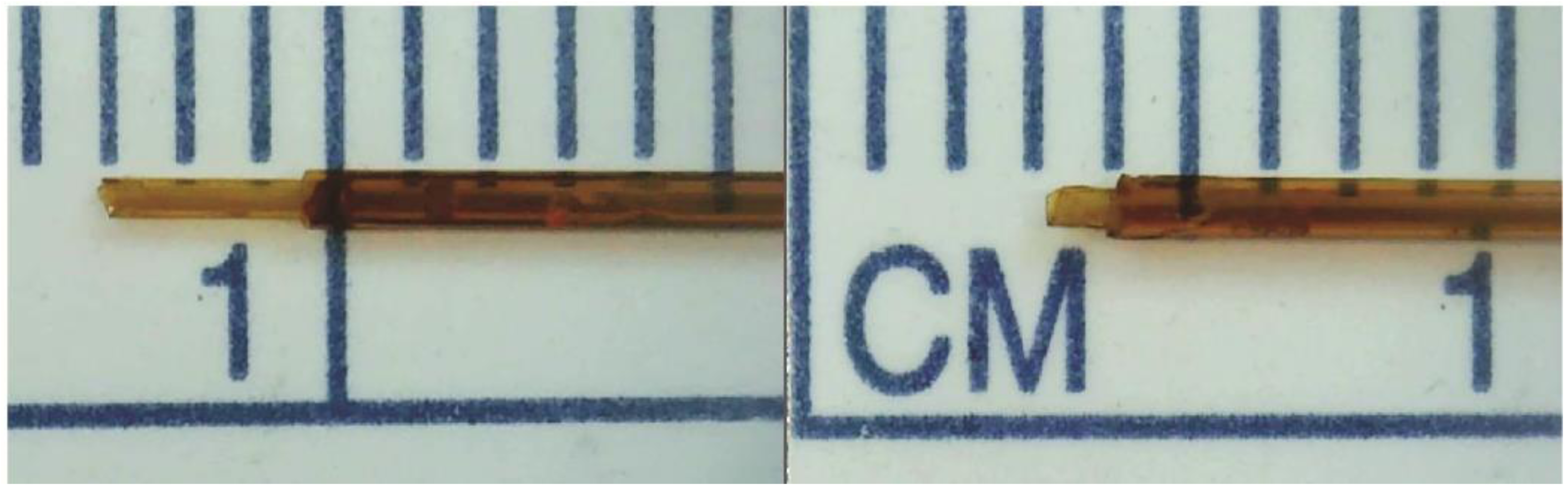
3-mm (left) and 1-mm (right) stepped-tip cannulas.

**Supplemental Figure 2.**
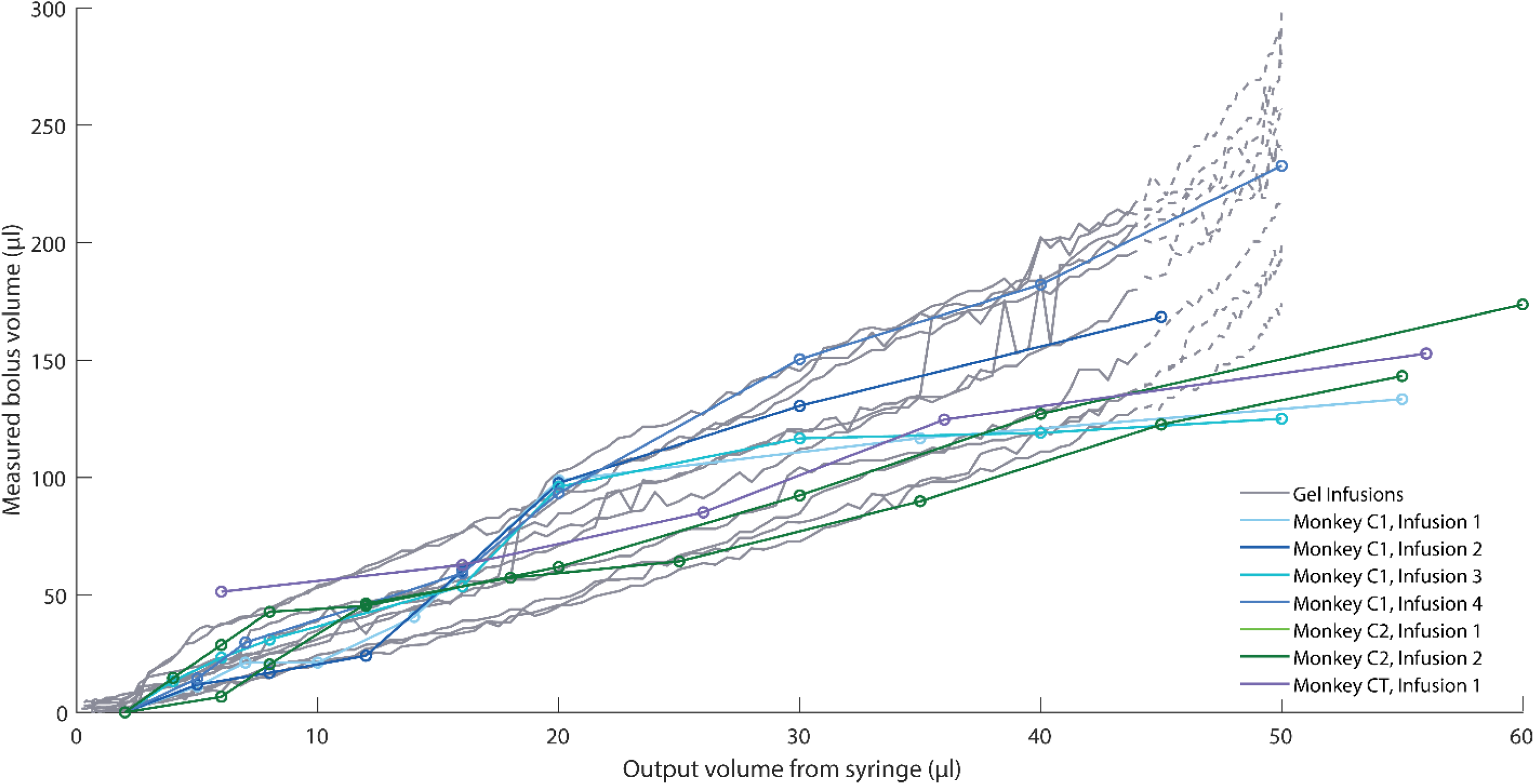
Comparison of agar and MRI cortical CED. This figure is the same as Figure 4B with the addition of data (dashed lines) from 44 µl to 50 µl which were omitted from statistical analysis due to their progressively steep upward trend which did not align with the MRI data. MRI data has been previously published [8].

**Supplemental Figure 3.**
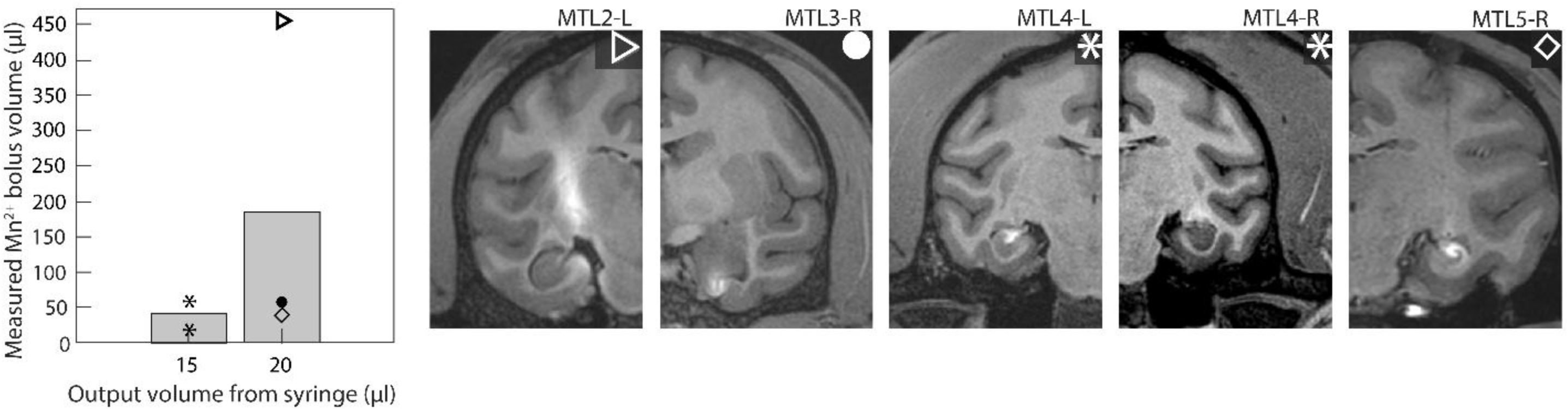
In addition to the deep infusions presented in Figure 6A, some infusions were not visually similar to cortical and thalamic data. These deep CED infusions were also made into the hippocampus, entorhinal cortex, and the tail of the caudate nucleus. Bar plots show the mean value of measured Mn^2+^ bolus seen in next-day, post-operative MRIs, with individual data points overlaid. Corresponding MRI slices in the coronal plane are shown. Shape labels correspond to each subject contributing to the data.

